# Reconfigurable multi-component nanostructures built from DNA origami voxels

**DOI:** 10.1101/2024.03.10.584331

**Authors:** Minh Tri Luu, Jonathan F. Berengut, Jasleen Kaur Daljit Singh, Kanako Coffi Dit Glieze, Matthew Turner, Karuna Skipper, Sreelakshmi Meppat, Hannah Fowler, William Close, Jonathan P.K. Doye, Ali Abbas, Shelley F.J. Wickham

## Abstract

In cells, proteins rapidly self-assemble into sophisticated nanomachines. Bio-inspired self-assembly approaches, such as DNA origami, have achieved complex 3D nanostructures and devices. However, current synthetic systems are limited by lack of structural diversity, low yields in hierarchical assembly, and challenges in reconfiguration. Here, we develop a modular system of DNA origami ‘voxels’ with programmable 3D connections. We demonstrate multifunctional pools of up to 12 unique voxels that can assemble into many shapes, prototyping 50 structures. Multi-step assembly pathways with sequential reduction in conformational freedom were then explored to increase yield. Voxels were first assembled into flexible chains and then folded into rigid structures, increasing yield 100-fold. Furthermore, programmable switching of local connections between flexible and rigid states achieved rapid and reversible reconfiguration of global structures. We envision that foldable chains of DNA origami voxels can be integrated with scalable assembly methods to achieve new levels of complexity in reconfigurable nanomaterials.

---

A key goal in molecular self-assembly is to build synthetic systems approaching the complexity of the molecular machines found in living cells, with emergent functions such as chemotaxis and adaptation.^1, 2^ The fundamental challenge is to achieve efficient self-assembly with high stability while also allowing for rapid transitions between different states and functions. Synthetic self-assembly has achieved remarkable advances, including supramolecular switches and materials,^3, 4^ *de novo* design of proteins,^5, 6^ and functional molecular devices and crystals made from DNA.^7–12^ However, many biologically inspired self-assembly strategies are yet to be fully explored. Biology exploits multifunctionality, from the pluripotency of stem cells at the macroscale to the reuse of protein motifs at the molecular level.^13^ Properties such as funnel-shaped energy landscapes, local-to-global folding processes, and the use of molecular chaperones have been proposed to explain the efficiency of native protein folding.^14, 15^ Similarly, recent theoretical models predict that local-to-global folding of linear chains could provide an effective strategy for molecular robotics.^16^ Here, we develop a system of modular DNA nanoscale ‘voxels’ to explore these approaches to increase yield and reconfiguration efficiency of multi-component nanostructures.

DNA is an excellent material for self-assembly of nanostructures due to its sequence-specific binding and ease of both synthesis and chemical modification. Diverse self-assembly principles have been demonstrated with DNA, including: algorithmic self-assembly,^17, 18^ hierarchical assembly,^19–22^ controlled nucleation of shapes,^23^ pathways,^24^ and micron-scale structures,^25^ allosteric shape change propagated across structures,^26^ paper-folding,^27^ molecular transport,^12^ self-limiting growth,^28, 29^ and crystal growth to macroscale structures with nanoscale features.^7, 11^

One of the most robust and frequently used approaches is the DNA origami method,^30, 31^ where a long single-stranded DNA (ssDNA) scaffold is folded into diverse nanostructures cross-linked by short ssDNA staple strands to form double-stranded DNA (dsDNA). DNA origami nanostructures provide a 3D molecular canvas or ‘pegboard’ of addressable pixels that allow for the spatial arrangement of guest molecules^11, 19, 20, 25, 32–35^ and reconfigurable interfaces for dynamic shape transformation.^8–12, 19–21, 25–27, 32, 33, 35–41^ However, scaling up DNA nanostructures in size and complexity is essential for advanced applications and is limited by the finite length of the scaffold strand (< ∼10,000 nucleotides (nt)) used in DNA origami, typically derived from the ssDNA genome of the M13 bacteriophage.

Hierarchical assembly of DNA origami subunits is a potential solution to scaffold size limits, but is currently hampered by low yields in 2 dimensions,^19, 27^ lack of rigidity in 3 dimensions,^21, 25, 41, 42^ the high cost of many unique DNA strands,^43^ and laborious processes to achieve new shapes such as redesign,^28^ reassembly,^25^ or stepwise remixing^19^ of components. Many current approaches also rely on fixed binding interactions between subunits with a high degree of stability,^19, 20, 25^ whereas reconfiguration typically requires flexible elements and reprogrammable interfaces.^27, 36–38^

Here, we present a set of 3-dimensional DNA nanostructure ‘voxels’ with internal and external connections that can be switched between inactive, rigid, and flexible states. DNA voxels are combined to form a multipurpose pool of components for assembly of diverse 2- and 3-dimensional hierarchical assemblies without redesign, refolding, or remixing of voxels. We then use the DNA voxels to explore stepwise assembly pathways involving folding of flexible DNA voxel chains, to improve assembly yield and achieve fast and reversible reconfiguration (Fig.1a).

## Construction of rigid DNA origami voxels

The hierarchical DNA origami assembly system described here builds on an existing DNA barrel design.^20^ The barrel is selected because it is rigid, high yield, does not form nonspecific aggregates though DNA blunt-end stacking, can polymerise along its coaxial axis, and can be used to template other materials on both inside and outside surfaces.^20, 32, 44, 45^ Here, we extend this by building a new monomer composed of two barrel subunits folded from the same scaffold and linked by a flexible scaffold tether. The monomer has 12 external interfaces for connections to tile 3D space (Fig. 1b). Each barrel forms an addressable ‘voxel’ in the 3D superstructure that can be specifically modified to host different molecular guests.

**Figure 1:**
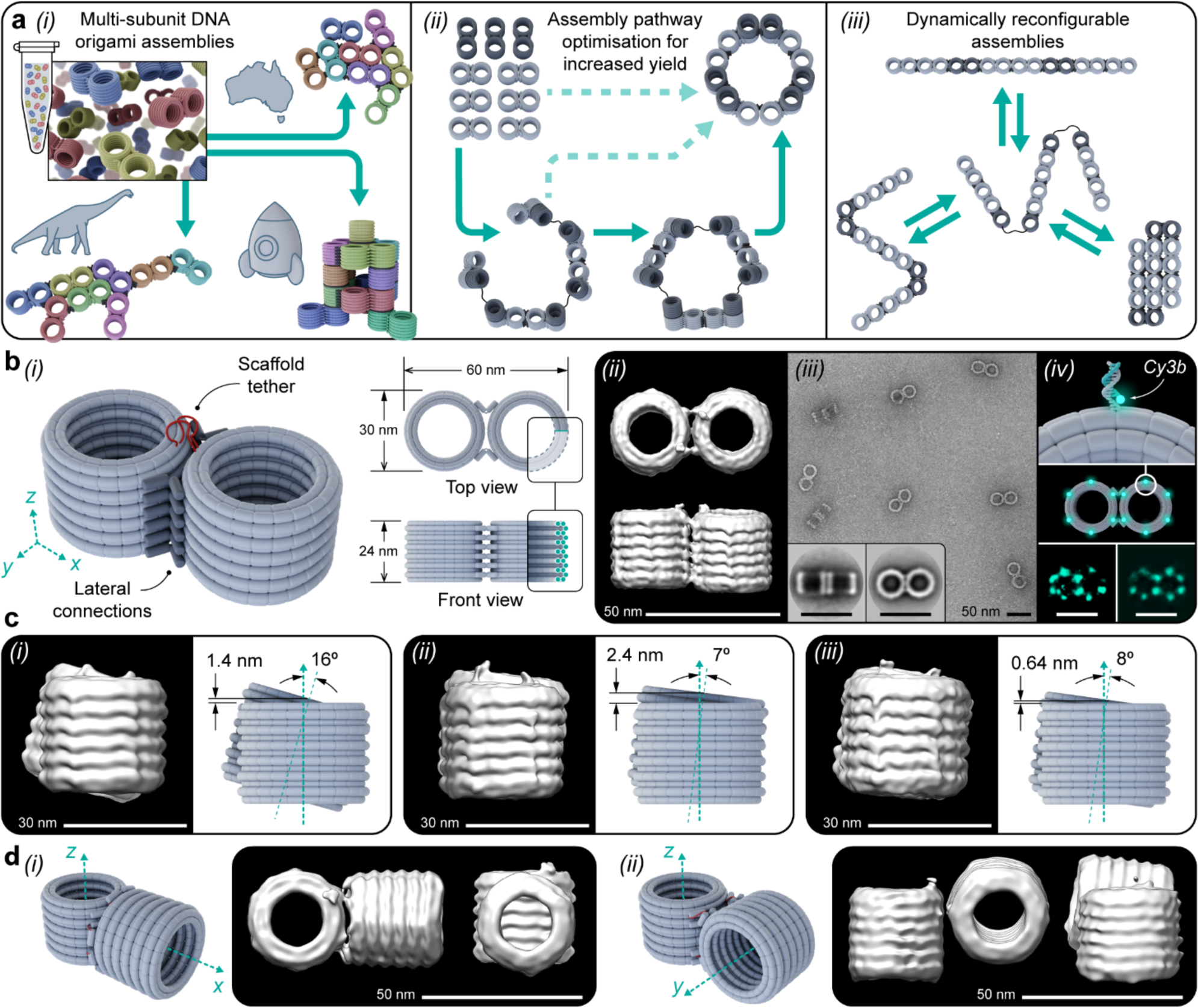
DNA origami monomers consist of two barrels, or ‘voxels’, linked together by rigid or flexible connections. **a** (i) Multiple shapes can be assembled from the same monomer pool by addition of different ssDNA connector strands, (ii) sequential addition of connector strands guides assembly pathways, (iii) toehold-mediated strand displacement of connector strands triggers dynamic reconfiguration. **b** (i) DNA origami monomer showing rigid intra-monomer connections (dark grey) and flexible scaffold tether (red); (ii) cryo-EM reconstruction (*N =* 3449); (iii) TEM and 2D class averages (*N_front_ =* 112, *N_top_ =* 166); (iv) DNA-PAINT reconstruction, top and middle: 12 docking strands hybridize fluorophore-labelled imager strands, bottom: example particle (left) and average (right, *N =* 10). **c** cryo-EM reconstructions (N = 7125, 3647, 3449, GSFSC resolution = 17.88Å, 17.83Å, 19.25Å respectively) show that (i) inward hinges result in 16° twist, (ii), horizontally alternating inward and outward hinges reduce twist but increase vertical offset to 2.4 nm, (iii) vertically alternating inward and outward hinges give lowest twist (8°) and offset (0.64 nm). **d** cryo-EM reconstruction of alternative intra-monomer interface designs with coaxial-axes in (i) z-x, and (ii) z-y directions (N = 1999, 3928, and GSFSC resolution = 18.54Å, 23.5Å respectively). Scale bars in **c** 30 nm, all others 50 nm.

Intra-monomer lateral connections between barrels are formed by ssDNA ‘connector’ strands that hybridize to ssDNA ‘plugs’ extending laterally from the side of each barrel, crosslinking the two barrels (Fig. 1b, Supplementary Figs 1,2). The interface consists of 24 7-nt plugs arranged in 2 vertical arrays with ∼15 nm spacing (Supplementary Fig. 3). Connectors consist of domains complementary to plugs on adjacent barrels separated by a 4-thymine (T) spacer or ‘hinge’. All 24 plug sequences are unique, and each interface has 12 unique 32-nt connector strands.

Negative-stain Transmission Electron Microscopy (TEM), cryo-Electron Microscopy Single Particle Analysis (cryo-EM), and DNA points accumulation for imaging in nanoscale topography (DNA-PAINT) were used to confirm the monomer structure (Fig 1b; Supplementary Figs 4-10, Supplementary Table. 1). TEM 2D class averages give monomer dimensions of 28 ± 1 (width) by 56 ± 1.5 (length) by 19 ± 3 (height) nm, in good agreement with design dimensions of 30 x 60 x 24 nm estimated using caDNAno software (helix width 2 nm, 0.34 nm/base pair).^46^ Measured spacing of docking sites in DNA-PAINT particle averages is 15 ± 0.7 nm (N = 5), in agreement with design spacing of 15 nm.

Cryo-EM reconstructions provide insight into the interface geometry (Fig. 1b(ii) and Supplementary Fig. 6-8, gold-standard Fourier shell correlation (GSFSC) resolution 17.83 - 19.25 Å). Unintended translational and rotational shifts were observed between the 2 subunits of the monomer (Fig. 1c, Supplementary Fig. 6-8 and 11). Coarse-grained simulations using oxDNA^47, 48^ predict that connector strands will form straight helices instead of bent hinges (Supplementary Fig. 12), leading to global monomer twist. Increasing the poly-T hinge predicted reduced twist but increased angular spread in oxDNA simulations, which would likely result in less rigid monomers (Supplementary Fig. 13-17 and Supplementary Table. 1). Instead, alternating patterns of inward and outward pointing hinges were tested experimentally. Vertically alternating hinge directions were found to result in the smallest twist and vertical shift (Fig. 1c(iii), Supplementary Note 1 and Supplementary Fig. 8). Alternative monomer geometries were also developed where the coaxial axes of the barrel subunits point in different directions, z-y, and z-x (Fig. 1d, Supplementary Fig. 18-22).

## Multi-component lateral and coaxial assembly of voxels

Voxels are assembled *via* inter-monomer lateral and coaxial connections (Fig. 2a, b). Unique monomers (Fig 1a. colours) are defined by 96 unique lateral plug sequences and 72 unique coaxial plug sequences (1212 nt), while core staple and miniscaf sequences are repeated (8952 nt, 88%). This approach allows for efficient reuse of most middle-inner staples and all miniscaf strands in all monomers (4848 nt, 49%). Voxels are folded in separate reactions, allowing for fully addressable modification of both interface and core sequences during monomer folding. Lateral connections are based on the intra-monomer design (Fig. 2a, Supplementary Fig. 23), while coaxial connections are based on previously published designs^20^ (Fig. 2b, Supplementary Fig. 24).

**Figure 2:**
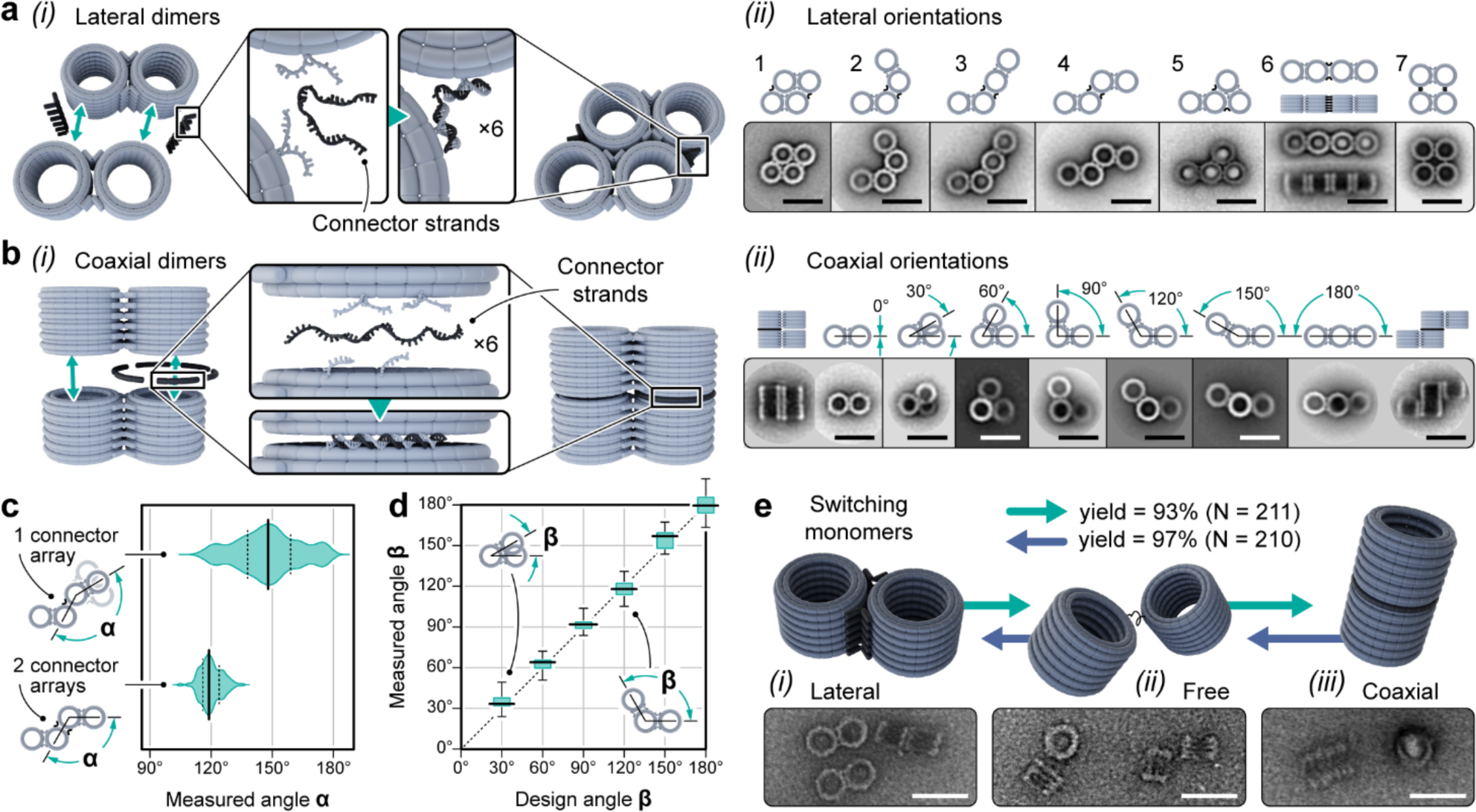
Programmable rigid connections between monomers. **a** (i) Lateral connections in *x-y* are formed by hybridization of connector strands to 7-nt plugs extending from outer helix staples, (ii) 7 unique lateral dimers. Top row: 2D models, bottom row: TEM class averages (left to right: *N =*39, 90, 37, 110, 24, 72, 30, 89). **b** (i) Coaxial connections in z are formed by connector strands hybridising to plugs extending from middle helix staples, (ii) 7 unique coaxial dimers. Top row: 2D models, bottom row: TEM class averages (from left to right: N =58, 29, 29, 30, 95, 303, 218, 257, 88). **c** Comparison of lateral dimer designs with one or two vertical arrays of connectors, the distribution of angles is smaller for the two-array design and the one-array design has a larger preferred angle. Angles measured from individual particles in TEM images (*N =* 120, 103). **d** Comparison of design and measured angle of the coaxial dimers in (b), angles measured from TEM particles (left to right: *N =* 60, 37, 41, 31, 18, 26). **e** DNA strand displacement triggers reversible switching of monomers between lateral and coaxial states, with forward and reverse yields of 93% and 97% respectively. Connector strands have a 7-nt toehold, a one-step process involves simultaneous addition of invader strands to remove existing connectors and new connector strands. Yields are calculated from TEM images. All scale bars 50 nm.

Connector strands were designed to successfully assemble all 14 uniquely identifiable dimers (Supplementary Note 1). Gel analysis was used to determine dimer band yield relative to the whole lane, then TEM was used to evaluate geometry and rigidity (Fig. 2a, b, Supplementary Figs. 25-54). All 7 lateral dimers had high yields (81% to 91%), while yields for the 7 coaxial dimers were slightly lower (45% to 86%) and depended on the connector sequence set (Supplementary Fig. 38). Using 2 coaxial connection sets instead of 1 increased yield, for example from 47% to 78% for the lowest yield connector set (Supplementary Fig. 55). Only the target dimer geometry was observed in TEM images for all designs indicating specificity of barrel connections. Reference-free 2D class averages were generated for TEM images using automated particle selection with no particles discarded, resulting in a small number of classes (n = 1-3). The fine structural detail visible in class averages shows particle uniformity and lack of deformation in dry state, providing evidence of rigidity (Fig. 2a). In some cases, blurred regions of class averages indicate flexibility or instability (e.g. lateral dimer 5, Supplementary Fig. 31).

A detailed analysis of the flexibility of lateral interfaces was performed for the ‘wave’ (Dimer 4), comparing one and two vertical connector arrays. The angular spread (Δα, s.d.) was found to reduce 3-fold from 15.4° for one array, to 5.2° for two, indicating an increase in rigidity (Fig. 2c). The single array had a higher average angle (148°), indicating that when flexible the hinge prefers to swing ‘closed’, likely caused by a combination of local blunt-end stacking within connectors and steric repulsion between barrels (Supplementary Fig. 13). The precision of rotational shift was also tested for the 7 coaxial dimers, in 30° rotation steps (β) from 0° to 180° (Fig. 2d). The observed rotation agreed with designed values, with low angular spread (Δβ =1.1° - 9°) indicating rigidity.

To enable switching by strand displacement, intra-monomer connector strands were designed with additional unique 7-nt toehold domains. ‘Invader’ strands, complementary to both toehold and connector domains, are added to displace connectors by toehold mediated strand displacement (TMSD)^49^. Monomers are then in a ‘free’ state where subunits can reorient but remain tethered allowing new structures to form on additions of alternative connector sets (Supplementary Figs. 56-61). Invader and alternative connector strands were added simultaneously in one step and switching yields were high (93 - 97%, 0.5 nM, Fig. 2e and Supplementary Figs. 59, and 60). However, at higher monomer concentrations (1 nM), the switching yield decreased to 75% due to increased dimerization (25%, Supplementary Fig. 61).

## Multifunctional monomer pools for 2D and 3D superstructures

A key goal of our design approach is to exploit the multifunctionality of a single set of monomers to rapidly prototype many different shapes. Starting with 3 unique monomers (Fig. 3a), monomers were initially folded separately then combined into a pool with equal concentrations and purified. The purified pool was divided into sub-pools and different connector strand sets were added to successfully construct 17 different trimers (Fig. 3b). Class averages show accurate shapes with good uniformity (n = 1-4 classes), while assembly yields were high (gel yield, average 86.1 ± 3.7 %, Fig. 3b and Supplementary Fig. 62). TEM images show only the correct geometry forms (Supplementary Figs. 63 - 96). We found that minimising the monomer twist (Fig. 1c) was essential to achieving accurate assembly of larger structures (Supplementary Note 1).

**Figure 3:**
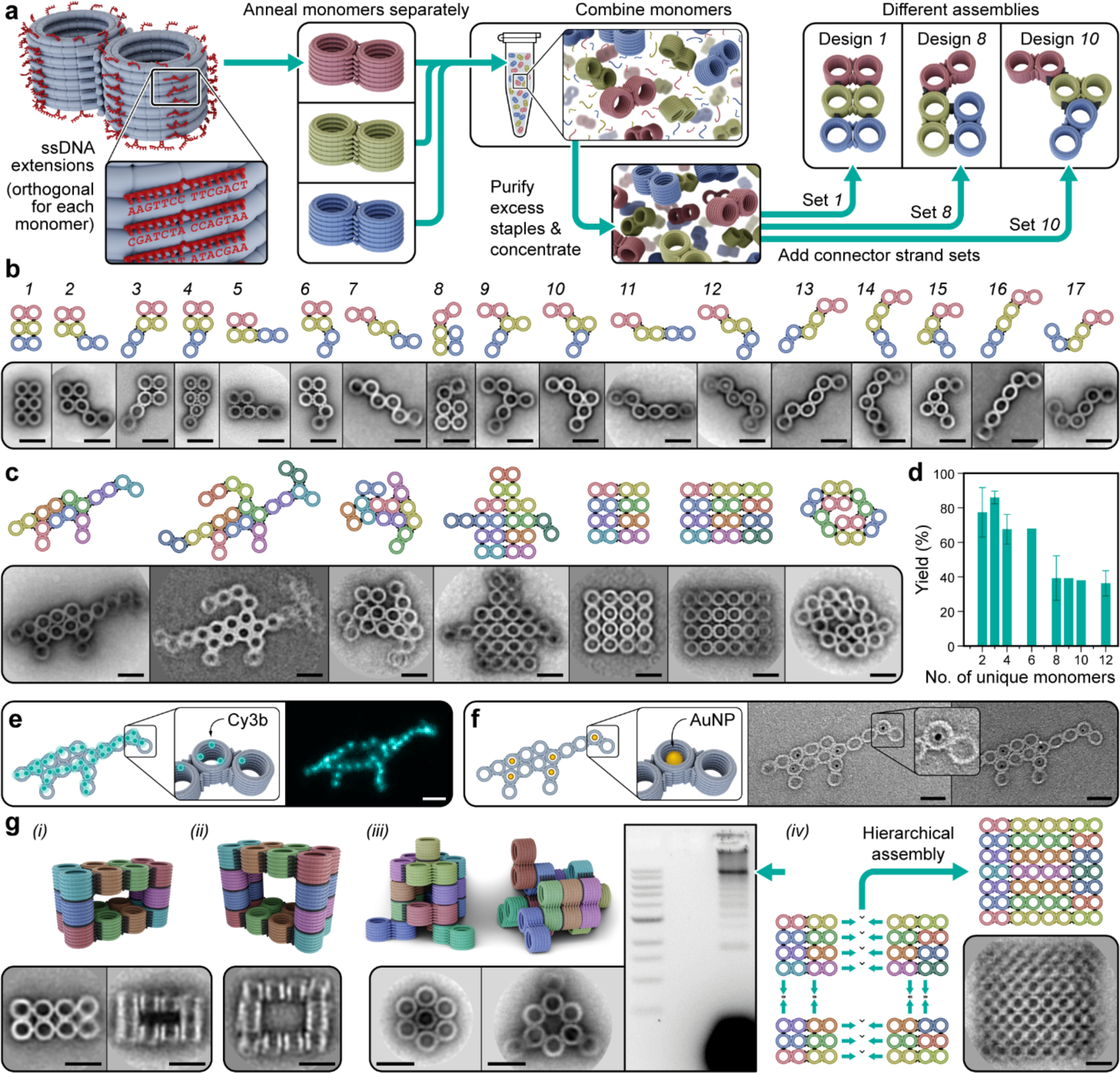
Multifunctional monomer pools assemble diverse 2D and 3D shapes. **a** Workflow to build different shapes from the same set or ‘pool’ of monomers. Left to right: unique monomers have unique plug sequences and reuse core sequences, are folded separately, combined, and purified as a monomer pool. The specific connector strand sets added determine the final structure. **b** Seventeen trimer assemblies built from the same trimer pool in (a), top row: rendered model and bottom row: TEM class averages with *N (left to right) =* 133, 74, 73, 82, 31, 99, 48, 51, 21, 38, 32, 32, 43, 38, 46, 36, 34, and 37. **c** Models and class averages of larger assemblies built from the 8 or 12 monomer pools with *N (left to right) =* 21, 11, 11, 11, 22, 21, and 21. **d** Assembly yield decreases with number of unique monomers, *N (number of designs) =* 14, 18, 3, 1, 3, 1, and 6, respectively. **e** DNA-PAINT of the dinosaur shape. Left to right: fully addressable pattern of DNA PAINT docking handles, and particle average (*N =* 10). **f** Gold nanoparticle patterning. Left to right: addressable pattern of gold nanoparticles, and TEM images. Selected monomers are folded with ssDNA handles on the inner surface to hybridise with ssDNA-conjugated 10 nm gold nanoparticles. **g** Scale up of assembly complexity and size. Model and class average for: (i) Small-cavity 3D box (*N =* 27), and (ii) Large-cavity 3D box (*N =* 10). (iii) A 12-mer 3D shape resembles a rocket. TEM class averages of the tip (*N =* 16) and base (*N =* 33) segments, and agarose gel electrophoresis result for entire structure showing optimised assembly yield of 12-mers. (iv) Fractal assembly process and class average of 28-mer rectangle (*N =* 25). In the first stage, monomers were assembled into 8-mer and 6-mer square and rectangles, respectively, then combined to form a fully addressable 28-mer rectangle, 210 x 240 x 30 nm. Scale bars 50 nm.

To demonstrate increased complexity, pools of up to 12 unique monomers were used to construct arbitrary 2D shapes, including 3 8-mers: dinosaur, Australia, rectangle, and 3 12-mers pool: dragon, robot, square (Fig. 3c, Supplementary Note 1, Supplementary Figs. 99 - 112). It was also possible to form 8-mer shapes, such as the dinosaur, from a 12-mer pool. TEM images confirm particles have the correct geometry. Generally, resolution of class averages and hence rigidity was observed to be higher for regions with more adjacent monomers (Fig. 3c). For example, monomers in the dinosaur body are connected to 3 adjacent monomers and have high resolution in the class average, indicating rigidity. The head monomer connects to 1 adjacent monomer and has lower resolution, indicating flexibility. In agreement with previous studies^19^, a sharp decrease in yield of 78 ± 14 % to 36 ± 7% was observed from 3- to 12-mers (Fig. 3d). Addressability of voxels in the resulting hierarchical assembly was confirmed using DNA-PAINT (Fig. 3e and Supplementary Fig 113). Lastly, we demonstrated a potential templating application by successfully patterning 10 nm gold nanoparticles inside specific voxels (Fig. 3f and Supplementary Fig 114).

Multifunctional voxel pools were then used to build more complex 3D shapes. Hollow 3D ‘boxes’ demonstrate cavity size control, a useful feature for templating nanoparticle assembly. Class averages of small and large box show cavity sizes with potential to capture spherical 20 and 40 nm nanoparticles, respectively (Fig. 3g(i,ii), Supplementary Fig 115-120). A 3D rocket ship was designed on a hexagonal lattice to test greater asymmetry (Fig. 3g(iii), and Supplementary Fig 121 - 123). Several design iterations were explored to optimize the connection pattern and resulted in an increase in yield from 5% to 33% of the 12-mer (gel yield, Supplementary Fig. 124). However, while class averages for subcomponent 4- and 8-mer rings show the correct structures by TEM (Fig. 3g(iii), and Supplementary Fig 125-127), the complete 12-mer collapsed under dry TEM conditions (Fig. 3g(iii)). Fractal, or multi-step, assembly strategies previously developed ^19^ were also used to construct a 28-mer rectangle from 2 × 8-mer and 2 × 6-mer sub-units (Fig. 3g(iv) and Supplementary Fig 128-129) and a ‘swirl’ by homo-dimerization of 4-mers (Supplementary Fig. 111).

## Improving assembly yield by folding monomer chains

Our results (Fig. 3d) and others^19^ show larger multi-component origami structures have decreasing yield (Supplementary Note 2), limiting potential applications. However, for a given monomer number some designs had higher yields (Fig. 3d), and monomer reconfiguration had high yields (93 - 97%, Fig. 2e). This suggests assembly paths with efficient transitions between high-yield intermediate or ‘precursor’ structures may improve yield.

A 9-mer 3D circle (Fig. 4a(i), structure ‘4’) was selected for pathway optimisation because single-step yield was very low (gel yield < 0.4%, Fig. 4a(iii)), with no correct circles found in TEM images. Instead, images of single-step products show polymer spirals (Fig. 4a(ii)). The single-array interfaces used in the circle have a preferred angle of 150 ± 15° in dimers (Fig. 2c), but here are required to form a 120° angle, potentially leading to strain that inhibits ring closing (Supplementary Fig. 130 - 132). Insight from supramolecular chemistry suggests that high-dilution conditions would shift assembly from polymerisation to cyclisation, however low concentration also reduces the yield of inter-monomer assembly. Thus, a flexible linear ‘chain’ was selected as a high-yield precursor, with higher yield double-array inter-monomer connections and flexible intra-monomer tether connections (Fig. 4a, structure ‘2’, Supplementary Fig. 133 - 135).

**Figure 4:**
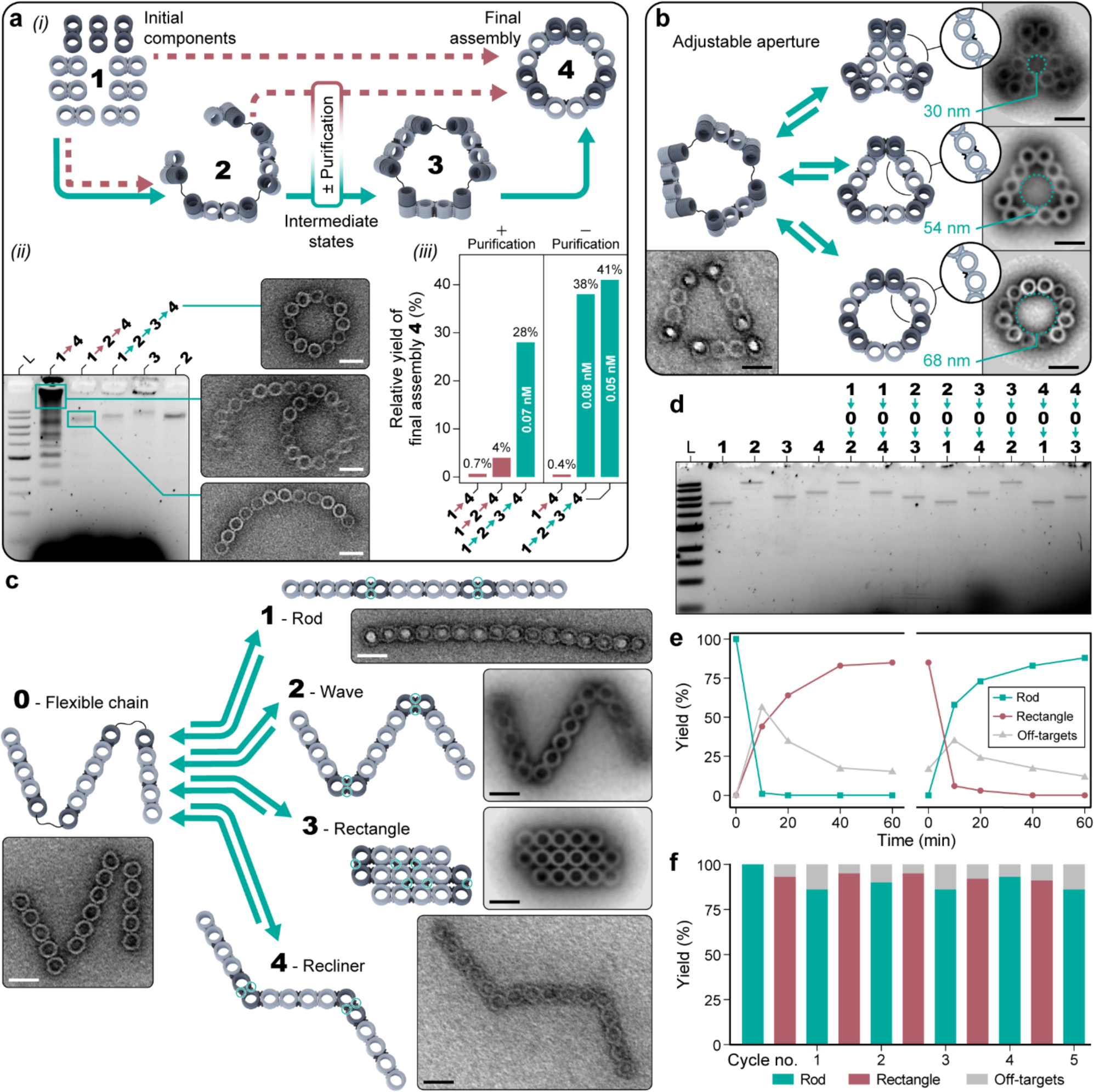
Programable reconfiguration of DNA origami voxel chains. **a** Optimization of assembly yield by directing assembly pathway, left to right: (i) schematic of voxel chain folding, (ii) agarose gel analysis of different pathways (left). Only path 1-2-3-4 (lane 4) results in high yield of the correct closed circle shape as determined by TEM. TEM images of mis-folded products from path 1-4 (lane 2) which results in chains or spirals, and path 1-2-4 (lane 3) which results in open shapes (right). (iii) Optimal assembly yield (unpurified chains, Supplementary Note 4) from monomers to the final shape is 100-fold higher for the path with the most sequential steps (1-2-3-4). **b** The 9-mer DNA voxel ring from (a) can also be reconfigured into 3 different shapes with central aperture diameter from 30 to 68 nm. Left: TEM image of the flexible ring (state 3 from panel a), Right, top to bottom: TEM class average of compact triangle, equilateral triangle, and circle (*N =*23, 90, 24). **c** An 8-mer flexible voxel chain can be switched between extended and compact states. Left: TEM image of the flexible chain (state 0, N = 1), Right top to bottom: TEM image of linear rod (state 1, N = 1), wave (state 2, *N =* 30), rectangle (state 3, *N =* 111), and recliner (state 4, N = 1). To reconfigure reversibly between states (blue arrows), invader and connector strands can be added to disrupt key interfaces (blue circles) and form the new structure. **d** Agarose gel analysis of switching reactions for pairwise transitions between states for the flexible chain. Invaders and connectors are added sequentially in this example so that sequences can be reused. **e** Switching yield as a function of time for the rod-to-rectangle pathway (i) and reverse (ii) from TEM images. In this example invaders and connectors are added simultaneously as no sequences are being reused. Rate constants can be estimated as 1×10^7^ M^-1^s^-1^ and 2×10^7^ M^-1^s^-1^ for forward (i) and reverse (ii) transitions, respectively, assuming second order reactions. **f** Switching yield for 10 consecutive steps (5 cycles) from TEM statistics (*N_average_* _=_ 147). Scale bars 50 nm.

The flexible linear chain was first assembled at high monomer concentration (1 nM per monomer), resulting in high yield (52 ± 5%, Fig. 4a(ii)). Unpurified or gel-purified chains were then folded into circles at low concentration (0.05 – 0.08 nM). While assembly yield increased 6-fold, it remained low (4%, gel-purified chain). TEM images show that polymerisation was avoided, but most circles failed to close. Thus, a further intermediate step of a flexible cyclized triangle was added (Fig. 4a(ii), structure ‘3’), in which the stronger double array interfaces are proposed to template ring closure. Finally, the flexible intra-monomer tethers were then switched to rigid connections and the double-array inter-monomer connections were removed to form single-array connections (Fig. 4a(i) and Supplementary Fig. 136). Remarkably, compared to a single-step the longest pathway involving 2 intermediate states resulted in up to 100-fold increase (unpurified chains, 0.4% to 41%, gel yield) with 40-fold increase for purified chains (0.7 % to 28%, TEM yield, excluding yield of purification step, Supplementary Note 4, Fig. 4a(iii), Supplementary Fig. 137-143, Supplementary Table 2).

## Dynamic reconfiguration of 2D and 3D voxel assemblies

Folding of monomer chains was then explored for dynamic reconfiguration of 2D and 3D shapes. Similarly to dimers (Fig, 2e), intra-monomer interfaces on ‘vertex’ voxels were switched between flexible and rigid states by addition of invader or connector strands. The length of the ssDNA scaffold tether limits the possible monomer geometries (Supplementary Fig. 144). The 269-nt tether has maximum ssDNA contour length of ∼180 nm, but in oxDNA simulations the tether has secondary structure (Supplementary Fig. 11). Experimentally, structures with tether spacings of 7 – 33 nm formed with high yields (N_designs_ = 7, 79 – 89%, Supplementary Fig. 144 and 146), but not for 62 nm, indicating an upper limit to the tether spacing (Supplementary Fig. 144).

Local to global reconfiguration was then tested for the 9-mer 3D circle to achieve reversible collapse and expansion of the central aperture. The flexible cyclised structure (structure ‘3’, Fig 4a) was used as a transitional structure, and vertex monomers (dark grey in Fig. 4b, Supplementary Fig. 147) were switched between 0°, 60° and 180°. At the same time, the inter-monomer interfaces were switched between concave, straight and convex states (Fig. 4b, inset). This resulted in reconfiguration between the compact triangle, equilateral triangle, and circle (Fig. 4b). Correct switching was confirmed by TEM for 1-step transitions (flexible-to-rigid and rigid-to-flexible), and for an example 2-step reconfiguration pathway (flexible-compact-flexible-circle) (Supplementary Figs. 148 - 159).

An 8-mer flexible chain with 2 vertex voxels was used demonstrate reconfiguration between 4 shapes with high yield (average 85 ± 5%, Fig. 4c and 4d, Supplementary Fig. 160-167). The switching yield for the rod-rectangle transition was measured as function of time and ∼80% yield was reached in 60 minutes (Fig. 4e, Supplementary Tables 3-4), a comparable timescale to previous origami multimer switching results^39^. Over 5 complete cycles between the rod (30 x 480 nm) and rectangle (90 x 180 nm), consisting of 10 total transitions, the yield of the target shape was consistently above 75% (Fig. 4f, Supplementary Tables 5-6), demonstrating repeatable high-yield reconfiguration at larger length scales.

## Conclusions

While custom multi-component DNA origami assemblies can be designed with high precision for specific functions, such as rotary molecular motors^10^ or multimaterial lattices^11^, our multipurpose voxel pools allow for more rapid prototyping of diverse shapes for new applications. Many applications require optimisation of geometric parameters such as cavity size, packing geometry and rigidity, at sizes greater than achieved by a single M13 scaffold (10 – 100 nm). For example, to build DNA-templated inorganic crystals with dynamic optical properties requires optimisation of cavity size to host nanoparticles > 50 nm for strong plasmonic effects and with periodicity on the order of the wavelength of light (200 – 800 nm) for photonic applications. Another important application area for multi-component DNA origami systems is as scaffolds for biophysical studies of large-scale protein assemblies approaching cell size (1-10 μm), such as chemotaxis arrays and the cytoskeleton. To mimic and study these biological systems, the DNA scaffolds must combine high yield assembly with rapid and efficient dynamic reconfiguration. Here we have shown that small local changes in geometry can achieve rapid changes in global structure on timescales suitable for driving protein reconfiguration in single-molecule biophysical experiments.

The foldable DNA voxel chain also provides a new system to rapidly explore a range of self-assembly strategies inspired by protein folding and supramolecular chemistry. We show that assembly pathways with high-yield intermediate states were able to rescue a 3D circle design from essentially zero yield to a usable yield. This system has future applications in exploring how strategies such as sequential reduction in conformational freedom, templating and reuse of functional motifs can be useful in multi-component origami assembly. In particular, longer foldable voxel chains could be folded into complex shapes, analogous to the way the ssDNA scaffold can be folded into DNA origami shapes. DNA voxel chains can also be used to explore how 3D origami chain displacement can mimic DNA strand displacement, similar to 1D nanorod^21^ or 2D tile displacement,^37^ to incorporate dynamic computing.

Overall, we have described multipurpose pools of DNA origami voxels that serve to rapidly prototype dynamic 3D multi-component DNA origami nanostructures. We envision that foldable chains of DNA origami voxels can be integrated with scalable assembly methods^40^ to achieve new levels of complexity in reconfigurable nanomaterials.^32, 33, 41, 50^ The voxels retain a regular arrangement of addressable surface sites for positioning of nanoparticle and protein guests with high precision. Interfaces are optimised for both high-yield, high-stability assembly, and rapid dynamic reconfiguration, and could be adapted to respond to more varied inputs including pH, small molecules and external electric and magnetic fields. Finally, foldable DNA voxel chains provide new methods to increase yield by optimizing assembly pathways, opening up future applications in optical and magnetic nanomaterials, single molecule biophysics, and synthetic cells.

## Materials and Methods

### Method 1: Folding of DNA origami voxels

DNA strands were purchased from IDT (Integrated DNA Technology) and resuspended in MilliQ water to minimum concentration 200 µM. Monomers were folded from 7308-nt ssDNA M13 scaffold (Guild BioSciences) and 10x minimum staple excess. Each 120 µL reaction consists of: 12 µL of scaffold (0.1 µM), 12 µL of core staples (1 µM), 12 µL of modified staples (1 µM), and 12 µL of connector strands (1 µM) with 12 µL of 10xTE (100 mM Tris-HCl, 10 mM EDTA, pH 8.0), 12 µL of MgCl_2_ (120 mM, final concentration 12 mM), and 48 µL of MilliQ water. After adding all the components, samples were vortexed and annealed in a PCR machine (Eppendorf Mastercycler Nexus) by heating to 65°C for 15 mins and then linearly cooling from 50°C to 40°C for 18 hrs.

### Method 2: Gel electrophoresis of DNA voxels

Agarose gel electrophoresis (Thermo Fisher Scientific, EasyCast Mini Gel Electrophoresis System) was used to separate correctly assembled monomers from the folding mixture. The gel contained 1.95 g of agarose powder (Bioline, final concentration 1.3% w/v) in 150 mL of 0.5 x TBE buffer (50 mM Tris, 50 mM Boric acid, 1 mM EDTA, 11 mM MgCl_2_, pH 8.3) and 3 µL of SYBR safe (Thermo Fisher Scientific). Gels were run at 70 V for approximately 2-3 hrs at room temperature. Gel images were captured using a Bio-Rad ChemiDoc imaging system with either Cy3 or SYBR safe filter setting. Gel image analysis and band densitometry were carried out using Bio-Rad Image Lab software. After imaging, the bands of interest were excised on a blue light transilluminator (Fisher Biotec) and the nanostructure was extracted from the cut gel band using a spin filter (Bio-Rad, Freeze and squeeze column) at 18,000g 20 min, 4°C. The concentration of DNA was measured on a Nanodrop spectrophotometer (Thermo Fisher Scientific), and this was used to calculate concentration of nanostructure given total DNA molecular weight.

### Method 3: Voxel purification using PEG precipitation

PEG precipitation was used to concentrate gel-purified monomers to useful concentration ranges for making assemblies and to remove free DNA in solution to reduce background noise in TEM images. The Mg^2+^ in DNA samples was increased to 20 mM by adding an appropriate amount of 1 M MgCl_2_. Equal volume of the DNA sample and 2xPEG buffer (15% 8000 PEG (w/v), 5 mM Tris, 1 mM EDTA, and 505 mM NaCl) were then mixed. The mixture was briefly vortexed and then centrifuged at 16,000 g, 25°C, for 25 mins. The supernatant was carefully removed using a pipette followed by a second centrifugation at the same setting for 4 mins to allow removal of the residual supernatant. The pellet was resuspended in 15 µL of 1xTE, 10 mM MgCl_2_ for 3 hrs at 30°C on a shaker at 1000 rpm.

### Method 4: Voxel purification using ultracentrifugation

To purify monomers, a rate-zonal centrifugation protocol with a 15% to 45% (v/v) glycerol gradient was used^51^. A commercial Gradient Station (BIOCOMP) unit was used to prepare the glycerol gradient by gently rotating a compatible tube containing two layers of 15% glycerol followed by 45% (1xTE, 10 mM MgCl_2_) at 20 rpm, 85° for 1 minute. The glycerol gradient was spun in a Beckman SW55-Ti rotor at 55,000 rpm and 4°C for 45 minutes. After that, the gradient was fractionated into 100 µL fractions using an automated system of the Gradient Station equipped with UV detection at 260 nm. For purification of large assemblies, a longer tube compatible with the SW41 rotor was used for gradient preparation. The tube was gently rotated at 15 rpm, 81.5° for 2.5 minutes and then spun in a Beckman SW41-Ti rotor at 41,000 rpm and 4°C for 25 minutes. Collected fractions were loaded into an agarose gel to resolve the content and determine fractions with DNA. Before TEM, one round of PEG precipitation was performed on the fractions of interest to exchange the glycerol buffer for 1xTE, 10 mM MgCl_2_ buffer. A typical fluorescent trace during the fractionating process is shown in Supplementary Fig. 164.

### Method 5: Multi-component assembly

Unique monomers were folded in separate tubes. After that, relative folding yield was determined by agarose gel electrophoresis. The band intensity of an arbitrary monomer was used as a reference for normalization to obtain relative folding yields of all monomers in the gel. Based on this information, unpurified monomers were combined into one pool with an appropriate volume to give equal monomer concentrations. Monomer pools were gel purified, and PEG precipitated to obtain purified and concentrated monomer pools. A typical unoptimized yield from M13 scaffold to gel purified monomer pool was 6%. For assembly, monomer pools were mixed with intra and inter-monomer connector strands, 1xTE buffer, and 1 M MgCl_2_ to obtain a solution containing 1 nM per unique monomer, 200 nM connector strands, 1xTE and 20 mM Mg^2+^. Inter-monomer connector strands were added to assemble monomers while additional intra-monomer strands were added to ensure monomer structure remains rigid during this step. The assembly mixture was incubated in a PCR machine at 40°C for 18 hours. For some structures multiple monomer pools were used. For example, when building the Rocket ship assembly, two pools were used for purification, one for lateral monomers and one for the coaxial monomer, due to the very different monomer recovery yields in the PEG precipitation step for these two sub-pools.

### Method 6: Negative-stain transmission electron microscopy (TEM) and class averages

For TEM imaging a plasma-treated grid with formvar film and heavy carbon coating (Ted Pella EM grids, GSCU300CH-50) was used. A 5 µL sample was pipetted onto the grid and incubated for 1 to 10 minutes, depending on the concentration of the DNA nanostructures, to ensure that enough particles would be present on the grid. The excess sample was blotted off with a filter paper, and 10 µL of 1xTE was quickly applied to the grid and blotted again to prevent the grid from drying out. Subsequently, 10 µL of a 2% uranyl acetate solution in water was added to the grid, which was then blotted immediately on a filter paper. This was done by dragging the grid horizontally on the filter paper to gradually soak up the stain droplet. TEM images were acquired on a JEOL 1400 in bright-field mode at an accelerating voltage of 120 kV.

2D particle class averages were derived from TEM micrographs using the RELION software^52^, enabling the assessment of structural attributes such as flexible and rigid areas. Particle picking and 2D classification were performed using RELION software, with an automated particle picking algorithm (Laplacian-of-Gaussian, RELION 3.0.6) employed for small assemblies (monomer, dimer, and trimer) and manual picking for larger structures. The number of resulting non-zero classes in RELION 2D classification was low (n = 1-4), typically clearly showing distinct particle orientations (top, side, reflected). To give an unbiased assessment of particle homogeneity, no particles were discarded, and all class averages are shown in Supplementary Figures.Some larger shapes required uniform particle orientation on the TEM grid for determination of accurate 2D class averages. To achieve this, TEM images were manually rotated before being integrated into the RELION workflow. Class averages were not relied upon for yield calculations.

### Method 7: cryo-EM imaging and particle reconstruction

Cryo-electron microscopy (cryo-EM) single particle analysis was used to elucidate structural details of monomers. Sample conditions were 3 uL, 700 nM nanostructure, and 1xTE buffer with 6 mM MgCl_2_. The preparation of specimens was carried out using C-flat grids, which were subjected to plasma cleaning using a Gantan Solarus plasma cleaner to ensure optimal hydrophilicity and cleanliness. Subsequently, the grids were plunge-frozen using a Vitrobot Mark IV, ensuring rapid vitrification and minimal ice crystal formation. The prepared grids were then loaded into a Thermo Fisher Scientific Glacios Cryo-EM, equipped with a 200kV Field Emission Gun (FEG) source.

Data acquisition was conducted on a Falcon 2 Direct Electron Detector, with Thermo Scientific EPU software automating the data collection process. The magnification and electron dose were set for data set in Fig 1 c(ii) such that the pixel resolution is 1.1 Å per pixel, and electron dose is 40 electrons per Å^2^; for all other cryo-EM models, the pixel resolution is 0.86 Å per pixel, and electron dose is 50 electrons per Å^2^.

The subsequent image processing workflow was executed using CryoSPARC (v4) for particle picking, 2D classification, and 3D reconstruction. Initially, a small subset of the micrograph data set was selected for manual particle picking, followed by 2D classification, and 3D reconstruction. After that, the 3D model was used to generate 50 templates, showing possible orientations of particles in the ice. These preliminary templates were then utilized for automated particle picking from all micrographs. Picked points were inspected and filtered by changing the power score and Normalized Cross-Correlation (NCC) score to remove empty areas. The particles were then subjected to 2D classification, allowing for the refinement of particle groups based on structural similarities. These averages served as the foundation for the 3D reconstruction process, ultimately yielding detailed and informative 3D models of the monomers.

### Method 8: Super resolution fluorescent microscopy (DNA-PAINT)

A TIRF microscope (Oxford NanoImager) was used to perform DNA-PAINT imaging. To prepare the sample, a protocol from the literature^53^ was adopted to create a customized chamber with immobilized DNA nanostructures and mobilized imager strands containing a cy3b fluorophore. In line with the protocol, a DNA origami tile was used as a fiducial marker. The key experimental parameters included 0.25 nM DNA barrel monomer, 0.25 nM DNA origami tile, and 5 nM imager strand. Imaging parameters consisted of 70 mW laser power, 300 ms exposure time, in total internal reflection mode. The localization of fluorophore binding sites was analyzed using the DNA-PAINT software (Picasso), which processed 5,000-10,000 time-lapsed images. Typical parameters in Picasso included baseline (400), sensitivity (0.47), quantum efficiency (0.8), and pixel size (117 nm).

### Method 9: Switching monomers between coaxial and lateral state

Monomers were folded, and gel purified. The same staple pools were used to fold both coaxial and lateral structures, but with different intra connectors. To switch between the two structures, lateral invader and coaxial connector strands were added to the purified lateral monomer, and coaxial invader and lateral connectors to the purified coaxial monomer. Concentrations used for the switching reactions were: 0.5 nM monomer, 100 nM connectors, 100 nM invaders, and 20 mM Mg^2+^. Switching samples were incubated at 40°C for 18 hours before TEM analysis.

### Method 10: Multistep ‘fractal’ assembly of the ‘swirl’ and the 28-mer rectangle

The swirl was built using a 4-mer that self-dimerizes to form an 8-mer structure. In stage 1, purified and concentrated monomers were used to construct first-stage assemblies. In stage 2, the previous assemblies were gel purified and connector strands were added. The solution was annealed at 38°C for 18 hours. Here the concentration values used were: 0.01 nM tetramer, 200 nM connector strands (intra and inter), and 20 mM Mg^2+^. The 28-mer rectangle was assembled using two 8-mer and two 6- mer assemblies in stage 1. In stage 2, sub-assemblies were purified, combined and connector strands added. The solution was annealed from 40°C to 30°C over 18 hours. Concentration values used were: 0.065 nM per assembly, 40 nM connector strands (intra and inter), and 40 mM Mg^2+^.

### Method 11: DNA-PAINT imaging of Dinosaur assembly

The slide preparation protocol as above was modified to increase the nanostructure incubation time to 10 min due to low particle concentration. For one sample: 5 µL of the unpurified Dinosaur assembly mixture was added to 3.18 µL of 1.57 nM gel-purified DNA tile, 2 µL of 0.5% Tween-20, and 9.82 µL of 1xTE, 10 mM Mg^2+^. Imaging parameters were as above used to acquire 10,000 time-lapsed images.

### Method 12: Assembly of the circle, compact triangle, and equilateral triangle

In step 1, flexible chains were assembled from monomers, and either used unpurified or gel purified. In step 2, 200 nM connector strands were added to 0.05-0.008 nM flexible chain, in 20 mM Mg^2+^ and incubated the sample at 40°C for 18 hours. In step 3, 400 nM connectors were added to 0.034 nM flexible triangles and incubated at 37°C for 18 hours. Alternatively, compact triangles or equilateral triangles could be assembled from the flexible triangle by adding different sets of connector strands. To determine the yield for the first two steps (monomers to flexible strands, and flexible strands to flexible circles), agarose gel electrophoresis was used. For the final steps, TEM was used to determine the percentage of particles with the correct geometry. To reduce background signal in TEM micrographs, PEG precipitation was used to remove free DNA strands.

### Method 13: Switching between rod (state 1), wave (state 2) and rectangle (state 3)

To perform the switching reaction, the first state was assembled, and gel purified, then incubated with invader strands for 6 hours at 37°C. Typical conditions used were 0.05 nM initial structure, 200 nM invader strands, and 20 mM Mg^2+^. Second, connector strands were added and incubated for 6 hours at 37°C. Typical conditions used were 0.04 nM flexible strand, 200 nM connectors, and 20 mM Mg^2+^.

### Method 14: Time-resolved switching

Multiple aliquots of purified structures were prepared, and connector and invader strands were added to each aliquot at time intervals of 60, 40, 20, 10, and 0 minutes. The mixtures were incubated at 40°C at concentrations: 0.07 nM rod structure, 200 nM connector and invader strands, 20 mM Mg^2+^, in 25 µL aliquot. After incubation, aliquots were immediately placed in dry ice to stop assembly. To prepare the TEM grids, frozen samples were defrosted by adding 20 µL of 1xTE to the 25 µL aliquot and quickly mixing with a pipette. The percentage of correctly switched particles was determined from TEM images and used to calculate the reaction rate constant, assuming a second-order reaction.

### Method 15: Multiple switching cycles

Five complete switching cycles, equivalent to 10 switching steps, were performed for 18 hrs at 40°C for each step. Using the same batch of starting material, 10 x 20 uL aliquots were prepared. In each switching step, the same connector and invaders solution was added to all aliquots to trigger switching. In addition, one aliquot was reserved and stored at −20°C; and this aliquot was imaged using TEM to determine the switching yield for a particular switching step. It was necessary to add increasing amounts of connector and invader strands to achieve high switching yields (Supplementary Fig. 165). For each switching step, the required amount of connector strand was added to achieve a constant concentration ratio of connector: invader of 2.3. Connector concentration (nM) used for step 1 to 10 is: 50, 110, 215, 330, 512, 678, 917, 1082, 1309, and 1411. Invader concentration (nM) used for step 1 to 10 is: 50, 48, 93, 143, 223, 295, 400, 470, 570, and 614. Except for the 1^st^ step, all steps 2 to 10 had a constant connector to invader concentration ratio of 2.3.

### Method 16: Gold nanoparticle conjugation

Citrate-coated gold nanoparticles (10 nm) were purchased from Sigma-Aldrich. To prevent salt-induced precipitation, the gold nanoparticles were pre-stabilized with a strong ligand (4,4- (Phenylphosphinidene)bis(benzenesulfonic acid) dipotassium salt hydrate). For conjugation, dithiol DNA purchased from IDT was added in 300x excess with phosphine-coated gold. The salt concentration was raised to 100 mM by adding a 1 M NaCl solution. The final concentrations of gold and DNA were 0.11 mM and 36 mM, respectively. The mixture was incubated at 25°C for 18 hours and then purified and concentrated using an Amicon filter (50K) with three washes of 1xTE buffer. The concentrated solution was stored at 4°C. To combine the AuNP-DNA complex with the nanostructure, samples were mixed at a 1:20 molar ratio and incubated at 40°C for 18 hours.

## Acknowledgement

The authors acknowledge funding from: the Australian Research Council (ARC): DE180101635 (S.W) and DP210101892 (S.M.); Westpac Research Fellowship (S.W., K.C.D.G, M.T.L.); The University of Sydney, The University of Sydney Nano Institute (J.F.B., M.T.L., J.K.D.S), and the University of Sydney Physics Foundation (M.T.L.); Australian Postgraduate Award (M.T.L., J.K.D.S, M.T); Australian Department of Industry, Science, Energy and Resources AUSMURIV000001 (M.T.L). The authors acknowledge the facilities as well as the scientific and technical assistance of the Australian Microscopy & Microanalysis Research Facility (ammrf.org.au) node at the University of Sydney: Sydney Microscopy & Microanalysis, and Sydney Analytical, a core research facility at the University of Sydney.

